# Shared Neural Responses and Individualised Emotional Experience: An fMRI Study of Music Listening and Anhedonia

**DOI:** 10.64898/2026.09.01.747608

**Authors:** Ayse Nilay Kamaci, Yanxing Chen, Marie-Stephanie Cahart, Owen O’Daly

## Abstract

Anhedonia, the diminished capacity to experience pleasure, is a core symptom of depressive disorders and is linked to altered reward, salience, and affective processing. However, whether individual differences in anhedonia are associated with variance in neural responses during naturalistic emotional experiences remains unclear.

This study examined neural synchrony, subjective emotional similarity, and continuous anhedonia-related variation in 28 healthy young adults who listened to emotionally evocative classical music during fMRI. Participants listened to happy, sad, and neutral excerpts while providing continuous emotional ratings. Inter-subject correlation (ISC) analyses quantified voxelwise neural synchrony across the brain and within affective and reward-related regions of interest (ROIs). Continuous inter-subject representational similarity analysis (IS-RSA) tested whether pairwise neural synchrony varied with anhedonic depression symptoms, measured using the Mood and Anxiety Symptoms Questionnaire (MASQ-AD).

Robust ISC was observed during music listening, particularly in bilateral auditory cortices (Heschls gyrus) and sensorimotor regions. Pairwise neural synchrony showed limited correspondence with similarity in continuous emotional subjective ratings, with significant associations primarily observed in auditory and somatosensory regions rather than classical affective regions. Anhedonia IS-RSA showed no reliable associations between MASQ-AD scores and pairwise neural synchrony across targeted affective and reward-related ROIs.

These findings suggest that shared neural responses during music listening are strongest in perceptual and sensorimotor systems, whereas subjective emotional similarity and anhedonia-related variation showed limited correspondence with neural synchrony. This pattern highlights the value of continuous, naturalistic approaches for distinguishing shared stimulus-locked processing from more variable components of emotional experience.

## Introduction

Major depressive disorder (MDD) represents a leading cause of global disability and is characterised by persistent low mood, cognitive impairments, altered emotional functioning, and reduced quality of life (American Psychiatric Association, 2013; World Health Organization, 2023). Anhedonia, broadly defined as a diminished capacity to experience pleasure, is a core feature of MDD associated with greater illness severity, poorer treatment response, and increased risk of suicide (Pizzagalli, 2014; Treadway & Zald, 2011; Uher et al., 2011; Luca et al., 2024; Winer et al., 2014). Although anhedonia is clinically prominent in MDD, milder anhedonic traits are also observed outside diagnosed clinical populations (Ely et al., 2021). Examining this variation dimensionally may therefore help characterise how hedonic capacity differs across individuals before or beyond the presence of a categorical diagnosis.

Neurobiological accounts of anhedonia implicate distributed systems supporting reward valuation, salience detection, interoception, and affective evaluation. Striatal hypoactivation during reward processing, particularly during reward anticipation and reward responsiveness, is among the most consistent findings in anhedonia, suggesting altered reward prediction and motivational processing (Der-Avakian & Markou, 2012; Pizzagalli et al., 2009; Whitton et al., 2015). Within this system, the nucleus accumbens (NAc) contributes to reward and emotional processing through the integration of signals related to reward, salience, and affective significance (Daniel & Pollmann, 2014; Koelsch, 2014). Its activity reflects sensitivity to rewarding and emotionally relevant stimuli and has been shown to vary with anhedonic traits, even within healthy populations (Keller et al., 2013). Beyond striatal dysfunction, regions involved in affective salience, interoception, and appraisal, including the amygdala, insula, and anterior cingulate cortex (ACC), are also relevant to anhedonia and music-evoked affective processing (Avery et al., 2014; Koelsch, 2014; Phelps & LeDoux, 2005). The amygdala supports detection and evaluation of emotionally salient stimuli (Phelps & LeDoux, 2005), while disruption of insular function has been linked to impaired interoceptive integration and reduced emotional differentiation (Craig, 2009). The ACC contributes to conflict monitoring, autonomic modulation, and affective evaluation (Bush et al., 2000). Alterations across these regions suggest that anhedonia may involve disrupted integration of sensory signals with their affective and motivational significance, rather than impairment of a single discrete emotion or pleasure-related circuit (Lindquist et al., 2012).

As anhedonia may involve distributed and integrative changes in how sensory information is linked with affective and motivational significance, it may be especially important to study these processes in contexts that unfold over time and approximate real-world emotional experience. Concerns regarding the ecological validity of conventional studies of affective cues, such as facial expressions (Ekman & Friesen, 1971), have therefore driven interest in naturalistic stimuli, such as movie clips or music, which engage richer and temporally extended perceptual, emotional, and cognitive processing that better reflect how emotions unfold in real-world settings (Hamilton & Huth, 2018; Schaefer et al., 2010).

Music is particularly well-suited for studying hedonic and affective processes, as it reliably evokes diverse emotional responses across individuals while recruiting neural systems involved in everyday emotional processing, including the amygdala, hippocampus, nucleus accumbens (NAc), ACC, and orbitofrontal cortex (OFC) (Juslin & Västfjäll, 2008; Koelsch, 2014; Mitterschiffthaler et al., 2007). Music-evoked emotional responses vary substantially across individuals (Krumhansl, 1997; Sloboda & Juslin, 2001), and higher trait anhedonia has been linked to lower pleasantness ratings and reduced activation in mesolimbic and paralimbic reward networks during music listening (Keller et al., 2013). Activity in the ventral striatum, particularly the NAc, has been positively associated with music-evoked pleasure, while variation in anhedonic traits has been associated with altered reward-circuit responsivity even in healthy populations (Koelsch, 2014; Keller et al., 2013; Heller et al., 2009). Together, these findings suggest that music provides a useful naturalistic context for examining clinically relevant variation in hedonic experience across a non-clinical sample.

However, naturalistic fMRI responses are difficult to characterise using conventional event-based general linear models (GLMs) that rely on prespecified stimulus events or modelled features (Monti, 2011; Sonkusare et al., 2019; Nastase et al., 2021). Inter-Subject Correlation (ISC) analysis provides a complementary, stimulus-driven approach by quantifying the temporal synchrony of blood-oxygen level-dependent (BOLD) signals across individuals exposed to the same stimulus (Hasson et al., 2004). Synchronised regional neural activity, associated with high ISC levels, is commonly interpreted as reflecting shared stimulus-locked processing or engagement, although it does not establish the specific processes underlying that similarity (Nastase et al., 2019). When combined with continuous subjective emotional ratings, pairwise ISC can identify shared neural responses which are linked to similar reports of subjective experience (Sachs et al., 2020; Trost et al., 2015).

While previous research has linked anhedonia to altered mean neural activation and functional connectivity during reward and music processing (Pizzagalli et al., 2009; Whitton et al., 2015; Keller et al., 2013), it remains unclear whether people with similar levels of anhedonia exhibit more similar time-resolved neural responses to the same naturalistic emotional stimulus. ISC is well suited to this question because it directly quantifies pairwise similarity in stimulus-locked neural time courses. IS-RSA combines this pairwise ISC framework with representational similarity analysis (RSA) logic by comparing neural similarity matrices with behavioural models constructed from continuous anhedonia scores (Kriegeskorte et al., 2008; Finn et al., 2020; Chen et al., 2020), avoiding the loss of information associated with dichotomising participants such as with a median split of the data (MacCallum et al., 2002; Royston et al., 2005).

Here, we examined pre-existing fMRI data and continuous emotional ratings from a final analytic sample of 28 healthy young adults (Cahart et al., 2024) listening to emotionally evocative classical music excerpts (Mitterschiffthaler et al., 2007). The study had three aims. First, we characterised the spatial distribution of voxelwise neural ISC, expecting robust synchrony in auditory regions engaged consistently by the shared musical stimulus. Second, we tested whether pairwise neural ISC corresponded with pairwise similarity in continuous emotional ratings, hypothesising correspondence within key affective regions, particularly the amygdala, insula, NAc, and ACC. Third, we used IS-RSA to determine whether participants with more similar MASQ-AD scores showed more similar stimulus-locked neural responses, particularly within affective and reward-related regions. In secondary analyses, we also examined whether participant pairs jointly higher in anhedonia showed altered neural synchrony. These analyses focused on the amygdala, insula, and striatal subdivisions, while whole-brain analyses were treated as exploratory.

## Methods

Data for this study were obtained from Cahart et al. (2024). Full details regarding study design, participant recruitment, and data collection are described in the original publication. The analyses reported here were conducted independently for the current research aims.

### Participants

The original study recruited 39 neurotypical, right-handed adults between the ages of 18 and 30 years, all of whom provided written informed consent. Participants were in good physical health, had no history of neurological or psychiatric conditions, and met standard MRI safety criteria. Data from 8 participants previously excluded by Cahart et al. (2024) were also excluded from the present analysis. A further 3 participants were excluded following extended quality-control checks specific to the current study. These decisions were informed by group-level MRIQC outputs, including framewise displacement (FD), DVARS, and signal-to-noise ratio (SNR) (Esteban et al., 2017). The final analytic sample therefore comprised 28 participants (15 male, 13 female) with a mean age of 22.4 years (SD = 4.13 years). As this was a secondary analysis of an existing dataset, no a priori sample-size calculations were conducted for the present analyses.

All procedures were approved by the King’s College London Human Research Ethics Committee (reference number HR-19/20-18771) and conducted in accordance with the Declaration of Helsinki. In line with the policies of King’s College London’s Department of Neuroimaging, all scans were visually inspected by the research team and reviewed by a neuroradiologist to confirm the absence of clinically significant abnormalities.

### Stimuli and Procedures

During fMRI scanning, participants listened to 13 classical music excerpts previously validated to evoke happy, sad, or neutral emotional responses in healthy adults (Mitterschiffthaler et al., 2007). While listening, participants continuously rated how the music made them feel using a scale ranging from-6 (very sad) to +6 (very happy). Auditory stimuli were delivered via headphones, and emotional ratings were made using a two-button response box: pressing the left button moved the on-screen slider toward-6, while pressing the right button moved it toward +6. All equipment was tested prior to scanning to ensure optimal auditory delivery.

The musical excerpts were presented in a fixed order for all participants to support time-locked group analyses. Each piece was separated by a 2-second silent interval. The full run comprised 3 happy, 3 sad, and 7 neutral songs. One neutral song opened the run, and each emotional song (happy or sad) was followed by a neutral one. Song durations varied slightly to avoid abrupt cut-offs. Full details of the musical stimuli, including the full list of songs and composers, are provided in Cahart et al. (2024). The order of presentation, stimulus onset and offset times, and corresponding song types are summarised in Table 1.

**Table 1.** Order of presentation of each musical piece, start and end times, and valence-types (happy, sad, or neutral piece).

| **Order of Presentation** | **Start Time (s)** | **End Time (s)** | **Song Type** |
| --- | --- | --- | --- |
| 1 | 0 | 29 | Neutral |
| 2 | 31 | 65 | Sad |
| 3 | 67 | 109 | Neutral |
| 4 | 111 | 147 | Happy |
| 5 | 149 | 183 | Neutral |
| 6 | 185 | 225 | Sad |
| 7 | 227 | 259 | Neutral |
| 8 | 261 | 301 | Happy |
| 9 | 303 | 337 | Neutral |
| 10 | 339 | 381 | Sad |
| 11 | 383 | 425 | Neutral |
| 12 | 427 | 467 | Happy |
| 13 | 469 | 507 | Neutral |

### Self-Report Measures

Before scanning, participants completed a battery of self-report questionnaires assessing their familiarity with the musical stimuli, musical background, and affective traits. These included subscales of the Big Five Inventory (BFI; John & Srivastava, 1999) to measure neuroticism, the Rumination-Reflection Questionnaire (RRQ; Trapnell & Campbell, 1999) to assess rumination, the Perth Emotional Reactivity Scale (PERS-S; Preece et al., 2018) to evaluate emotional reactivity, and the anhedonic depression subscale of the MASQ (Watson et al., 1995). Although data were collected on all these measures as part of the broader protocol, the present analyses focused on anhedonic depressive symptoms measured using the MASQ-AD. MASQ-AD scores were treated as a continuous measure of anhedonia severity in all anhedonia-related analyses.

### MRI Data Acquisition

All participants were scanned using a 3-Tesla MR scanner (Discovery MR750, General Electric, Milwaukee, WI, USA) equipped with a 12-channel head coil at the Centre for Neuroimaging Sciences, Institute of Psychiatry, Psychology and Neuroscience, King’s College London. Structural T1-weighted images were acquired using a magnetisation-prepared rapid gradient-echo (MPRAGE) sequence with the following parameters: repetition time (TR) = 7.35 ms, echo time (TE) = 3.04 ms, flip angle = 11°, slice thickness = 1.2 mm, and in-plane resolution 1.05 mm. Functional images were acquired using a 2D multi-slice gradient-recalled echo-planar imaging (EPI) sequence with the following parameters: TR = 2 s, TE = 33 ms, flip angle = 75°, slice thickness = 3 mm, field of view = 240 mm, and matrix size = 64 × 64. At the start of each fMRI run, 4 dummy scans (8 seconds) were acquired to allow signal stabilisation, these “dummy scans” were not saved and were not included in the subsequent analyses. The full fMRI run lasted 8 minutes and 27 seconds.

### MRI Data Preprocessing

All neuroimaging data were preprocessed independently from the original study to ensure that quality control and analysis procedures were tailored to the present research aims. Raw imaging files were first converted to Brain Imaging Data Structure (BIDS) format (Gorgolewski et al., 2016).

MRI quality assessment was performed using MRIQC version 22.0.1 (Esteban et al., 2017). Functional and anatomical MRI data were preprocessed using fMRIPrep version 25.0.0 (Esteban et al., 2018). The fMRIPrep workflow included slice-timing correction, head-motion realignment, functional-anatomical co-registration, spatial normalisation to MNI152NLin2009cAsym space, brain extraction, tissue segmentation, and estimation of nuisance/confound variables. Motion correction was performed using MCFLIRT (FSL; Jenkinson et al., 2002). Functional images were co-registered to each participant’s anatomical T1-weighted image using FreeSurfer’s mri_coreg (Fischl, 2012) and FSL’s FLIRT (Jenkinson & Smith, 2001) with boundary-based registration (Greve & Fischl, 2009). Spatial normalisation was performed using nonlinear registration with Advanced Normalization Tools (ANTs; Avants et al., 2008). T1-weighted images underwent N4 bias-field correction (Tustison et al., 2010), brain extraction, and tissue segmentation using FAST (FSL; Zhang et al., 2001) to identify cerebrospinal fluid, white matter, and grey matter regions.

Additional denoising was performed using custom Python scripts to regress out motion parameters, framewise displacement, DVARS, and low-frequency drifts modelled using discrete cosine bases. Anatomical component-based noise correction (aCompCor; Behzadi et al., 2007) was applied by extracting principal components from eroded cerebrospinal fluid (CSF) and white matter (WM) masks. The residual BOLD time series were bandpass filtered between 0.009 and 0.08 Hz and spatially smoothed with a 6 mm full-width at half-maximum (FWHM) Gaussian kernel in Statistical Parametric Mapping 12 (SPM12; Friston et al., 2007).

### Region of Interest Preparation

ROI and mask-based analyses were conducted alongside exploratory whole-brain analyses. For ROI-level ISC and neural-rating correspondence analyses, a priori ROIs were selected based on prior literature implicating the insula, ACC, and NAc/limbic striatum in anhedonia, affective evaluation, interoception, and reward processing (Bush et al., 2000; Craig, 2009; Daniel & Pollmann, 2014; Der-Avakian & Markou, 2012; Treadway & Zald, 2011).

ROI masks for the insula and ACC were generated using the WFU PickAtlas (Maldjian et al., 2003; RRID:SCR_007378) tool in SPM12, while the NAc was defined using the limbic striatum mask from the Mawlawi ROI atlas (Mawlawi et al., 2001). These masks were used to extract ROI-level ISC summaries and to assess associations between neural ISC and subjective-rating similarity within each ROI. Figure 1 visualizes the lateralized ROI masks used for ISC analysis.

**Figure 1.**
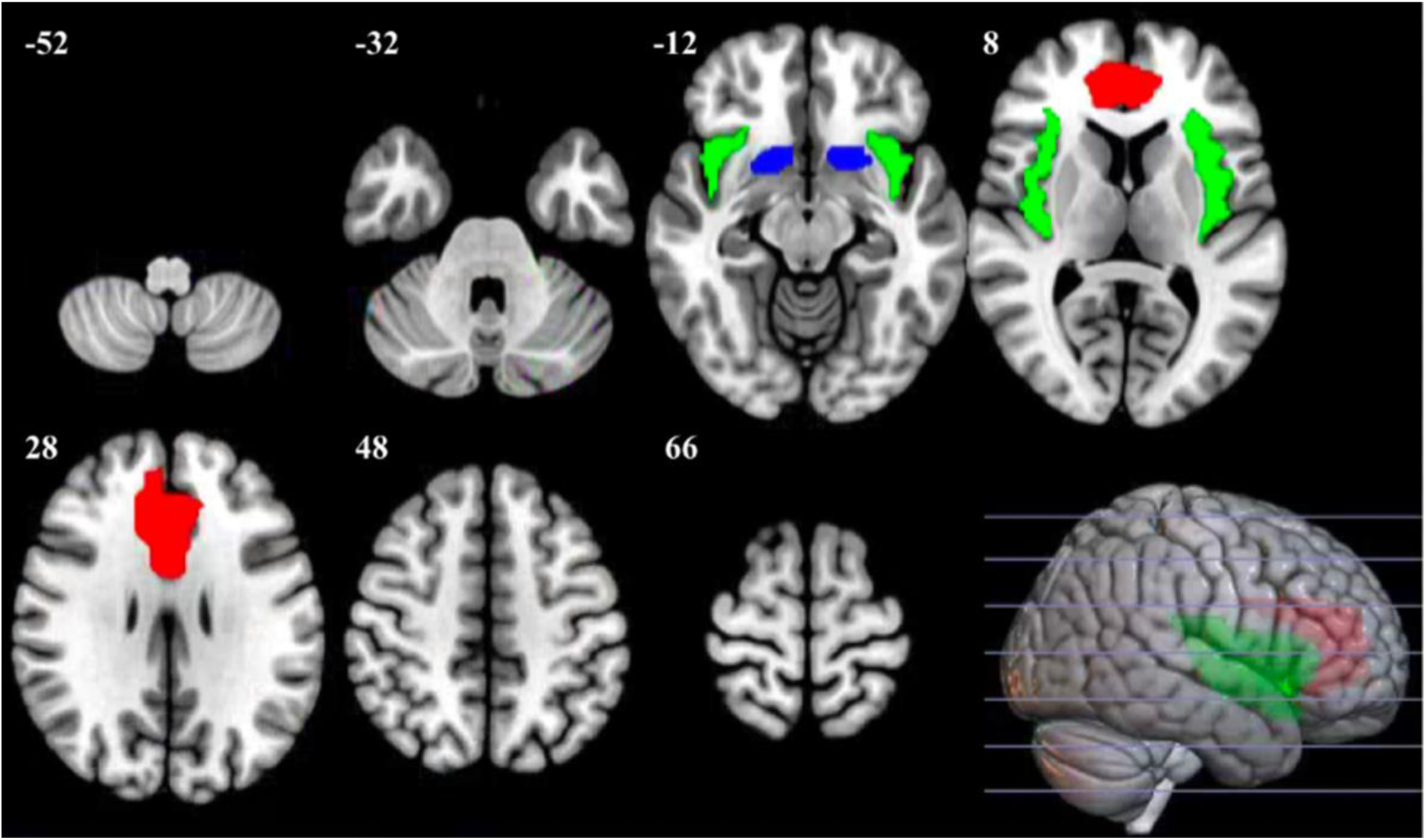
ROI masks used for ISC analysis. Brain regions of interest include the ACC (red), insula (green), and limbic striatum (blue) and are shown overlaid on axial brain slices (MNI z-coordinates indicated). A 3D rendering displays the spatial distribution of the ROIs on the lateral brain surface. Masks were visualised using MRIcroGL (https://www.nitrc.org/projects/mricrogl/).

For the anhedonia IS-RSA analyses, targeted masks comprised left and right amygdala, left and right insula, and left and right Mawlawi associative, limbic, and sensorimotor striatal masks. Exploratory analyses were additionally conducted using a whole-brain mask. Spatial alignment and resampling procedures for these masks are described below.

### ISC Analyses

Voxelwise ISC analyses quantified the similarity of BOLD time series across participants by computing pairwise Pearson correlation coefficients at each voxel (Hasson et al., 2004; Kauppi et al., 2014). Analyses were conducted using the ISC GUI Toolbox in MATLAB R2020a (The MathWorks Inc., 2020; Kauppi et al., 2014), producing inter-subject correlation matrices that represented neural synchrony across participant pairs during music listening.

Group-level ISC maps were visualised using the ISC Toolbox’s visGUI and SPM12. For ROI-level summaries, custom MATLAB scripts extracted values from the group-level ISC outputs within each bilateral ROI. Mean ISC values were computed for each ROI, and corresponding t-statistic values were extracted from the ISC statistical maps. These measures provided descriptive region-specific summaries of neural synchrony.

### ISC-Behaviour Correlation Analysis

Voxelwise correlation analyses were conducted to examine whether neural synchrony corresponded with similarity in subjective emotional responses. Continuous emotional ratings were correlated across all participant pairs to generate a behavioural ISC matrix. The upper triangle of this matrix was extracted and vectorised to form a behavioural ISC vector representing pairwise similarity in emotional ratings.

For each voxel within the whole-brain analysis mask or predefined ROI masks, pairwise ISC values were extracted across participant pairs to create neural ISC vectors in the same participant-pair order. Pearson correlation coefficients between the neural ISC vectors and the behavioural ISC vector were computed using a custom MATLAB script. This analysis was conducted voxelwise across the whole-brain analysis mask and separately within the predefined ROIs. No participant pairs were excluded from the behavioural ISC vector after final sample selection. The resulting correlation coefficients were mapped back into brain space and saved as NIfTI images for visualisation. Corresponding p-value maps were also generated and corrected for multiple comparisons; however, because pairwise observations are not fully independent, these analyses were interpreted cautiously as exploratory neural-behavioural correspondence analyses. Whole-brain ISC maps and neural-rating correspondence maps were thresholded using FDR correction at q < .05, with an additional minimum cluster size of 50 voxels used for cluster-level reporting.

### Anhedonia IS-RSA

IS-RSA was conducted to examine whether similarity in anhedonia severity was associated with pairwise neural synchrony. MASQ-AD scores were matched to the canonical participant ordering stored in the ISC Toolbox parameters using participant identifiers. All 28 participants had complete questionnaire data and were retained, producing 378 unique participant pairs.

For each ROI or exploratory mask, pairwise Pearson ISC coefficients were extracted from the existing whole-brain ISC Toolbox output for voxels falling within that mask. ISC was not rerun separately within each ROI. Extracted ISC coefficients were retained in the same upper-triangular participant-pair order as the behavioural matrices. Two behavioural model matrices were constructed from MASQ-AD scores. The primary anhedonia-similarity model was defined as the negative absolute pairwise difference in MASQ-AD scores, such that higher values represented greater similarity in anhedonia severity. As MASQ-AD was represented by a single continuous score, this reflected absolute univariate score difference rather than a multivariate distance metric such as Euclidean or Mahalanobis distance. The secondary Anna Karenina (AnnaK) model was defined as the mean rank of the MASQ-AD scores for each participant pair, such that higher values represented pairs with higher average anhedonia ranks and therefore tended to identify pairs jointly higher in anhedonia. Lower values represented pairs with lower average ranks; this model did not specifically measure disparity between pair members.

At each voxel, Spearman’s rank correlation was used to quantify the association between the pairwise neural ISC vector and each behavioural model vector. The primary anhedonia-similarity model and secondary AnnaK model scores were analysed separately. No covariates were included in the IS-RSA models. Statistical significance was assessed using subject-level permutation testing, with correction for multiple comparisons described below.

### Masks and Spatial Alignment

For the anhedonia IS-RSA analyses, targeted masks included left and right amygdala and insula masks, and left and right associative, limbic, and sensorimotor striatal subdivisions from the Mawlawi atlas. Exploratory analyses were also conducted using a whole-brain mask. Each mask was resampled into the ISC image space with nearest-neighbour interpolation, preserving its categorical definition. Resampled masks were binarised, visually checked against the ISC reference image, and saved to provide an exact record of the voxels included in each analysis.

### Permutation Inference

Statistical inference accounted for the non-independence of participant pairs using subject-level Mantel permutations (Mantel, 1967). For each of 5,000 permutations, participant identities were randomly permuted at the behavioural-matrix level before the upper triangle was re-extracted and correlated with the voxelwise neural ISC vectors. Pairwise entries were not permuted independently. All tests were two-sided, and permutation p-values were calculated as (b + 1)/(m + 1), where b is the number of permuted statistics at least as extreme as the observed statistic and m is the number of permutations. This prevents p-values of zero when using a finite number of random permutations.

Benjamini-Hochberg false discovery rate (FDR) correction (Benjamini & Hochberg, 1995) was applied across voxels within each mask, and statistical significance was defined as FDR-adjusted p < .05. These corrections were mask-specific and did not correct across the complete set of masks and models; consequently, within-mask corrected findings were treated as exploratory. Family-wise error (FWE) correction using the maximum absolute correlation across voxels within each mask was computed as a conservative sensitivity check but was not used as the primary inference criterion. For visualisation only, uncorrected permutation p-values were also transformed as –log10(p). This transformation did not alter statistical significance.

### Data and Code Availability

The dataset analysed in the present study was obtained from Cahart et al. (2024). In line with the original publication, the dataset is available from the corresponding author on reasonable request. Analysis scripts and derived result files generated for the present study are available from the corresponding author on reasonable request.

## Results

### Participant Demographics and Behavioural Metrics

The final analytic sample included 28 healthy young adults (15 male, 13 female), aged 18 to 30 years (M = 22.4, SD = 4.13). MASQ-AD scores ranged from 14 to 54 (M = 34.89, SD = 8.84), reflecting variability in anhedonia severity across the sample. MASQ-AD scores were retained as a continuous measure for the anhedonia IS-RSA analyses.

During the fMRI task, participants provided continuous emotional ratings on a scale from-6 (very sad) to +6 (very happy). Ratings corresponded with stimulus valence: happy excerpts elicited positive ratings (M = 3.38, SD = 1.53), sad excerpts elicited negative ratings (M = – 2.38; SD = 1.78), and neutral excerpts elicited ratings near zero (M = 0.46; SD = 2.35), indicating that ratings varied in the expected direction across stimulus valence categories. Figure 2 plots the time series of mean emotional ratings across the task duration.

**Figure 2.**
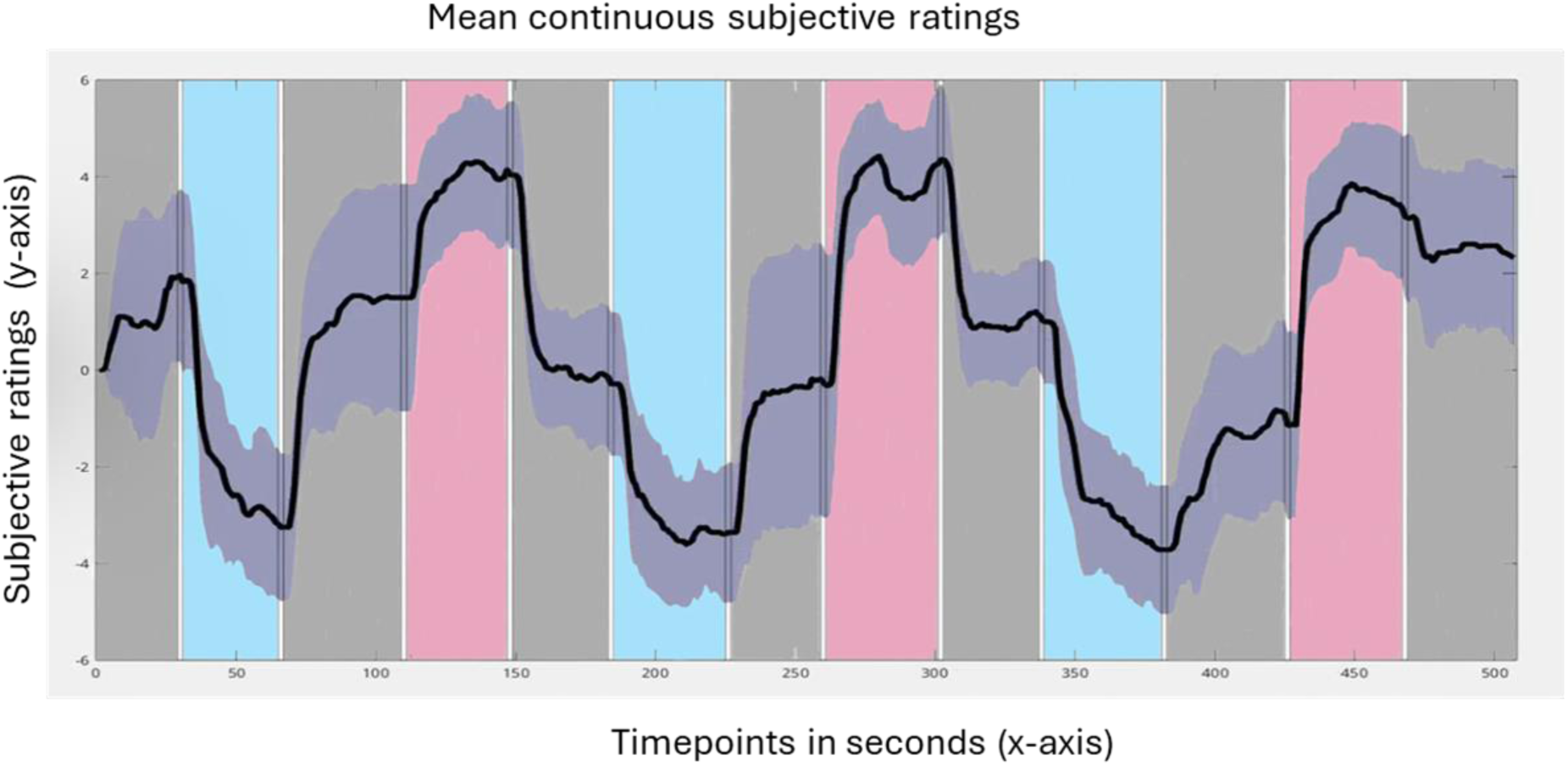
Mean continuous subjective ratings across all participants during the whole duration of the task. Emotional ratings range between –6 (very sad) and 6 (very happy). The black line represents the mean ratings throughout the task, and the surrounding purple areas represent the 95% confidence interval. Blue-coloured areas represent timepoints when sad songs were playing; pink-coloured areas represent timepoints when happy songs were playing; grey-coloured areas represent timepoints when neutral songs were playing. Plot was generated using custom MATLAB scripts (MathWorks, 2020).

### Inter-Subject Correlation Results

#### ROI-Based ISC

Mean ISC values (r̅) were calculated within the three a priori-defined ROIs used for the original ISC summary analysis. Positive synchrony was observed across all ROIs, with the highest mean ISC in the insula (r̅ = .0122), followed by the limbic striatum (r̅ = .0092) and ACC (r̅ = .0048). One-sample t-tests indicated significant synchrony in the insula (t(27) = 3.44, p = .002) and limbic striatum (t(27) = 2.78, p = .010), both of which survived Bonferroni correction across the three ROI tests (p < .017). Synchrony in the ACC was not significant (t(27) = 1.39, p = .18).

#### Whole-brain Voxelwise ISC

Voxelwise ISC analysis identified 42 clusters surviving FDR correction (*t*(27) > 4.08; minimum cluster size = 50 voxels). The strongest ISC was observed bilaterally in Heschl’s gyrus (left: *t*(27) = 68.29; right: *t*(27) = 59.62). Additional clusters emerged in sensorimotor and parietal areas, including the left postcentral gyrus, supplementary motor area, and inferior frontal gyrus. The largest cluster (19,961 voxels) spanned the parieto-occipital cortex and peaked in the right superior parietal lobule (*t*(27) = 14.45). Figure 3 depicts the voxel-wise ISC map across the brain.

**Figure 3.**
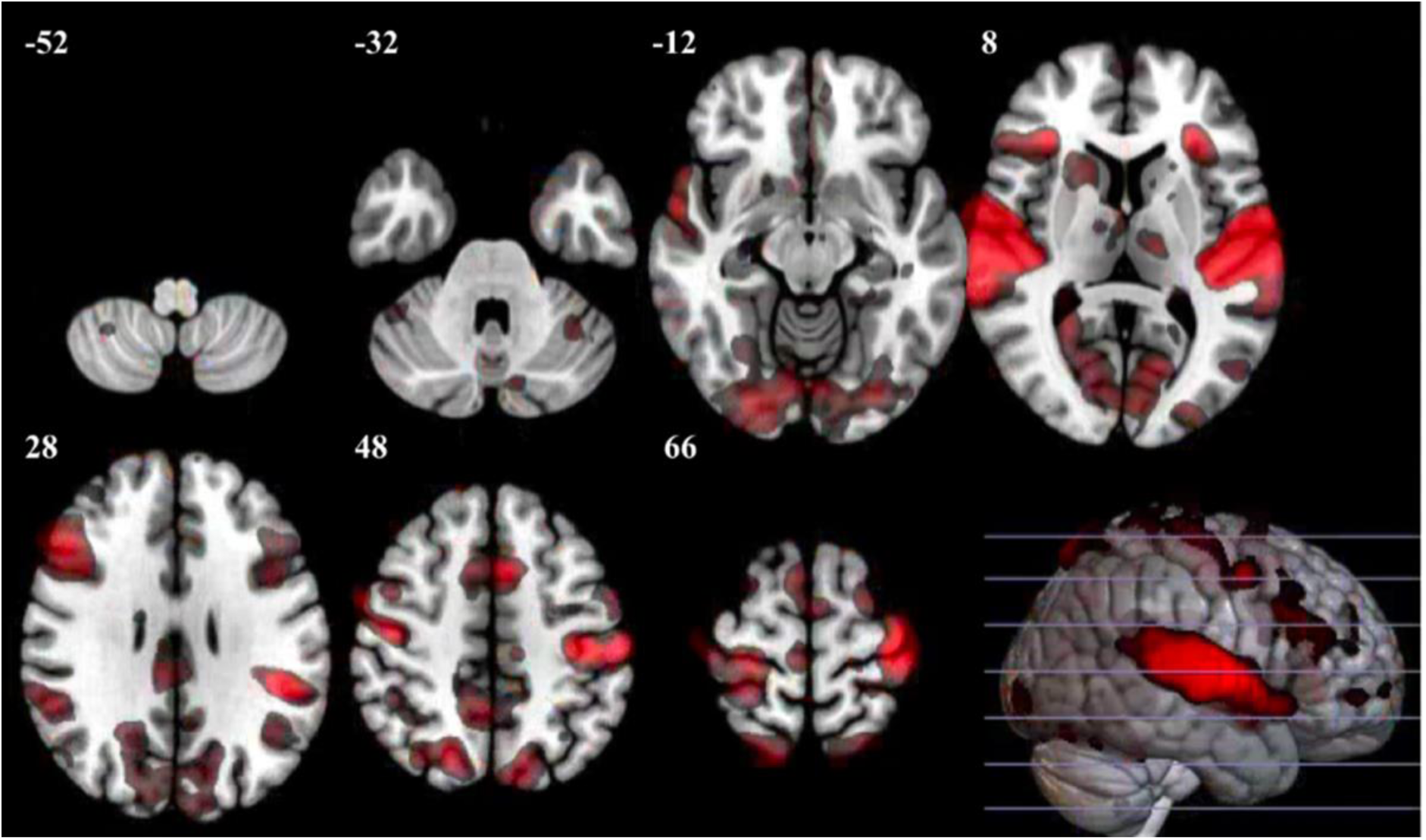
Voxel-wise ISC t-statistic map depicting significant neural synchrony during music listening. Results are thresholded at t > 4.08 (FDR-corrected) and overlaid on the SPM152 template using MRIcroGL (https://www.nitrc.org/projects/mricrogl/). Axial slices highlight peak effects in sensorimotor and bilateral auditory cortices (MNI z-coordinates indicated). Red hue brightness corresponds to increasing ISC strength, with brighter hues indicating stronger correlations.

### Neural Activity-Subjective Ratings Correlational Analysis Results

#### ROI-wise Neural-Behavioural Correspondence

Pearson correlations were computed between neural ISC and inter-subject similarity of emotional ratings within lateralised ROIs. Correlations were small, ranging from –.04 to .09, with the strongest effect occurring in the left limbic striatum (r = .091). No ROI-level correlation survived FDR correction.

#### Neural-Behavioural Correspondence Across the Whole-Brain

Exploratory voxelwise analyses identified 39 clusters showing associations between neural ISC and emotional-rating similarity surviving FDR correction (q < .01; cluster size ≥ 50 voxels). The largest cluster was located in the left postcentral gyrus (peak r = .271). Additional clusters were observed in the auditory and temporoparietal regions, including the right Heschl’s gyrus, right middle and superior temporal gyri, left superior temporal gyrus, and left supramarginal gyrus. Figure 4 ilustrates key regions showing significant ISC-rating correlations.

**Figure 4.**
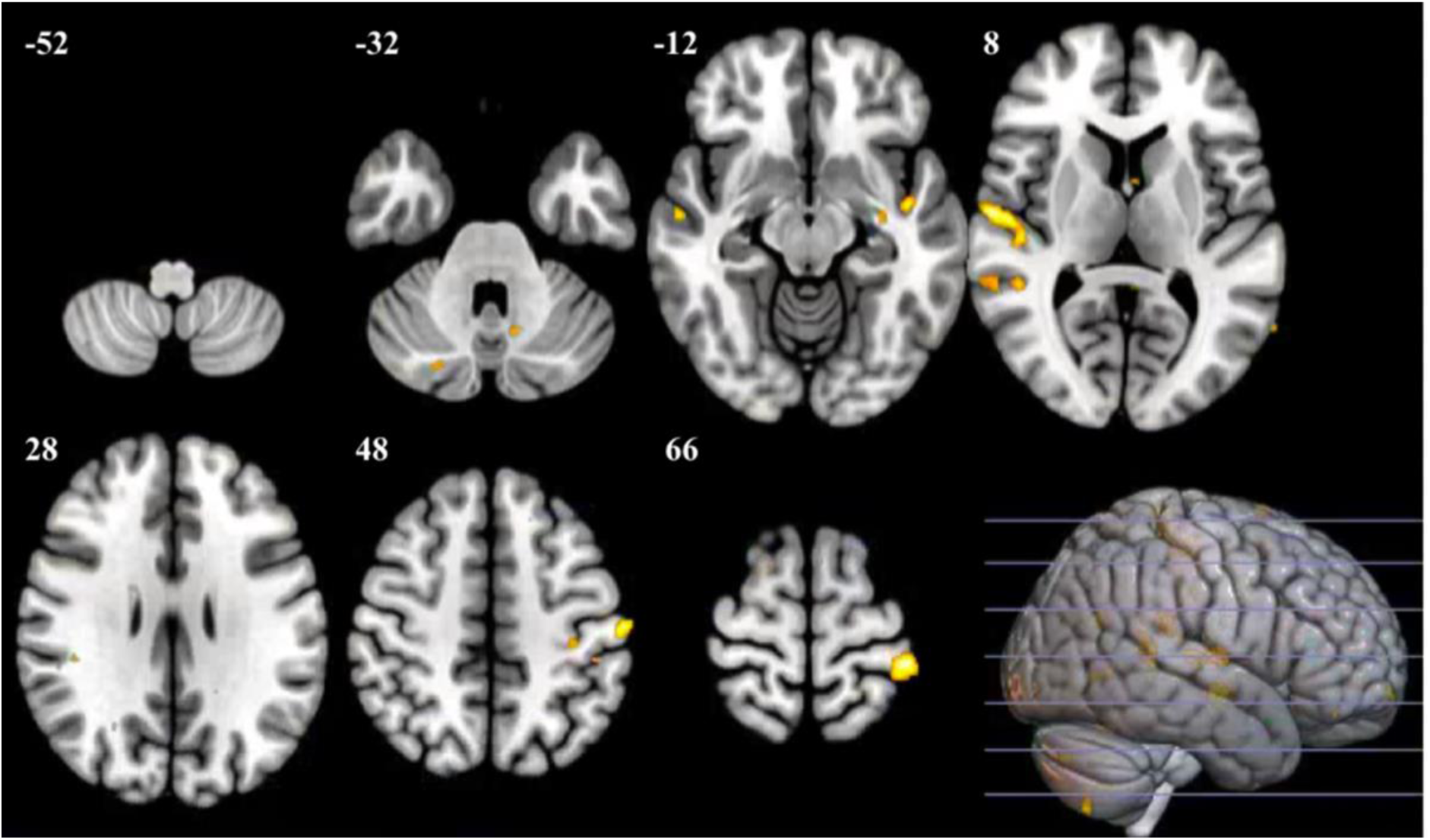
Voxel-wise map of significant correlations between neural ISC and emotional rating similarity. Clusters surviving FDR correction at p < 0.01 are overlaid on the SPM152 template using MRIcroGL (https://www.nitrc.org/projects/mricrogl/). Axial slices are presented, with the peak voxel displayed in the left postcentral gyrus (r = 0.271) (MNI z-coordinates indicated). Yellow hue brightness corresponds to increasing ISC strength, with brighter hues indicating stronger correlations.

### Anhedonia IS-RSA Results

IS-RSA analyses tested whether pairwise neural synchrony varied as a function of MASQ-AD scores using a primary anhedonia-similarity model and secondary AnnaK-based model. Results are reported numerically rather than as corrected statistical maps, as these analyses were hypothesis-led ROI analyses.

Across the targeted ROI analyses, neither the primary anhedonia-similarity model nor the AnnaK model showed reliable associations between pairwise neural synchrony and MASQ-AD scores. In the primary anhedonia-similarity model, a single right amygdala voxel showed a negative association surviving within-mask FDR correction (peak rho = –.187, p_unc = .0002, pFDR = .0496; peak MNI = 21.5, –8.5, –16.5), but this effect was spatially isolated and marginal and is reported for completeness rather than interpreted as evidence for a robust anhedonia-related effect.

The secondary AnnaK model did not show any effects surviving within-mask FDR correction. A focal negative left insula effect survived within-mask FWE correction only but did not survive FDR correction. This result was therefore treated as exploratory. No other targeted ROI or exploratory whole-brain analysis showed effects surviving FDR correction.

## Discussion

This study examined shared neural responses, subjective emotional similarity, and continuous anhedonia-related variation during naturalistic music listening. Robust ISC was observed across participants in bilateral auditory cortices and sensorimotor regions, with weaker synchrony in limbic and insular regions. Neural synchrony showed only modest correspondence with similarity in continuous emotional ratings, and these associations were primarily observed in auditory and somatosensory regions rather than classical affective ROIs. Continuous anhedonia IS-RSA provided limited evidence that MASQ-AD scores were systematically associated with pairwise neural synchrony. Most targeted ROI analyses and the exploratory whole-brain analyses showed no FDR-corrected effects. However, using the primary anhedonia-similarity model, a focal negative association in the right amygdala suggested that greater similarity in MASQ-AD scores was associated with lower neural synchrony in this region. Overall, these findings suggest that shared neural responses during music listening were strongest in perceptual and sensorimotor systems, whereas anhedonia-related variation in synchrony was limited and regionally focal within an affective salience region.

### Neural Synchrony During Music Listening

Robust synchrony in auditory cortices, sensorimotor areas, the precuneus, and superior parietal lobule is consistent with previous findings of widespread neural synchrony during naturalistic music listening and broader evidence for auditory, motor, and affective networks in music processing (Abrams et al., 2013; Koelsch, 2014; Zatorre & Salimpoor, 2013).

Auditory cortex synchrony likely reflects shared encoding of acoustic features such as pitch, rhythm, and timbre, which form the perceptual building blocks of musical experience (Zatorre & Salimpoor, 2013). Sensorimotor synchrony may reflect rhythmic entrainment or motor simulation during music perception (Chen et al., 2008; Zatorre et al., 2007). However, shared task demands, including button-pressing during continuous ratings, may have contributed to motor-related ISC, particularly around song transitions where rating changes were more likely (Nastase et al., 2019).

ISC was also observed in parietal regions, including the precuneus and superior parietal lobule. These effects may reflect shared attentional engagement or self-referential processing during music listening, although the present analysis cannot isolate the specific cognitive processes underlying synchrony in these regions (Cavanna & Trimble, 2006).

By contrast, classical emotion-processing regions exhibited weaker and more spatially restricted synchrony. The weaker synchrony observed in the insula may reflect greater inter-individual variability in emotional experience, despite its role in integrating sensory and affective information (Craig et al., 2009; Chang et al., 2013; Uddin et al., 2015). Significant ISC was not observed in the ACC, a region implicated in affective appraisal and regulation (Bush et al., 2000; Etkin et al., 2011). This pattern suggests that sensory and motor responses to music were more consistently shared across participants, consistent with their role in processing raw stimulus features. By contrast, evaluative or regulatory processes involved in constructing emotional meaning may vary more across individuals, consistent with evidence that higher-order association regions show greater inter-individual variability than unimodal sensory regions (Mueller et al., 2013).

Taken together, this spatial pattern suggests that the strongest ISC effects reflected common stimulus-locked processing of the musical excerpts and shared task engagement across auditory, sensorimotor, and attentional systems, rather than emotion-specific processing alone. However, ISC alone cannot determine which specific acoustic, motor, or cognitive processes drove synchrony in each region.

### Limited Correlation Between Neural and Emotional Rating Synchrony

The largest neural-rating correspondence cluster was located in the left postcentral gyrus. Given that participants used their right hand to provide continuous ratings, this effect may partly reflect shared somatosensory or motor aspects of the rating task, particularly if participants adjusted their ratings at similar moments, such as song transitions. One possible interpretation of the limited correlation, between neural and subjective ISC in brain regions linked to affective valuation, interoception, and emotion regulation (Craig, 2009; Etkin et al., 2011), is that music-evoked emotional experience depends on appraisal and meaning-making processes that vary substantially across individuals. Although happy and sad excerpts elicited ratings in the expected direction, variability was evident across participants, particularly for excerpts which were considered neutral relative to the affectively laden musical pieces. This is consistent with accounts of emotion proposing that affective experience is shaped by prior experience, expectation, context, and conceptual interpretation (Barrett & Bar, 2009; Lindquist et al., 2012; Seth & Friston, 2016). The relatively abstract and semantically sparse nature of music may amplify this variability, allowing listeners to construct emotional meaning through personal, cultural, and autobiographical associations (Juslin, 2013; Zatorre & Salimpoor, 2013). Consequently, shared neural responses to musical structure may only partially map onto shared subjective emotional experience, consistent with evidence that music-evoked affective and reward responses vary substantially across individuals (Sachs et al., 2016). In sum, these findings are consistent with the view that affective experiences are influenced by internal models, contextual expectations, and personal meaning-making.

### Anhedonia Severity and Neural Synchrony

Continuous IS-RSA provided limited evidence that anhedonia severity was systematically associated with pairwise neural synchrony during music listening. Across the targeted ROIs and exploratory whole-brain analysis, most effects did not survive within-mask FDR correction. In the primary anhedonia-similarity model, a focal negative association was observed in the right amygdala, suggesting that participants with more similar MASQ-AD scores showed lower neural synchrony in this region.

The secondary AnnaK model results did not survive the primary FDR correction. Although the exploratory left insula effect may be consistent with reduced coupling between shared subjective experience and shared neural responses among participants higher in anhedonia, the model did not directly test neural-behavioural correspondence. This interpretation therefore remains tentative and should be examined directly in future analyses.

Taken together, the results do not support a broad disruption of shared neural synchrony during naturalistic music listening in relation to anhedonia. Instead, they suggest that anhedonia-related variation, if present, may be focal and region-specific, particularly within affective salience and interoceptive regions such as the amygdala and insula (Phelps & LeDoux, 2005; Craig, 2009; Uddin et al., 2015).

### Strengths, Limitations, and Future Directions

This study has several methodological strengths. The use of naturalistic music listening improved ecological validity by capturing the temporal richness and complexity of real-world emotional experiences (Sonkusare et al., 2019). ISC further provided a stimulus-driven approach for identifying shared neural dynamics without relying on predefined response models, which is particularly useful for stimuli whose emotional features unfold continuously (Nastase et al., 2019). The integration of continuous emotional ratings enabled neural synchrony to be compared with pairwise similarity in subjective experience. Finally, the continuous IS-RSA approach retained individual variability in MASQ-AD scores and avoided dichotomising anhedonia into high– and low-symptom groups.

Despite these strengths, several limitations merit consideration. First, the modest sample size of 28 participants limits statistical power to detect subtle or higher-order effects, a common concern in neuroimaging research (Button et al., 2013). The sample was also non-clinical, young, and demographically homogeneous, which constrains generalisability to clinical populations or individuals with more severe anhedonia (Henrich et al., 2010). Although the use of a non-clinical sample reduced some confounds common in clinical cohorts, such as medication exposure, illness chronicity, and psychiatric comorbidity, it also limits generalisability to individuals with more severe anhedonia or diagnosed affective disorders. Larger and more demographically diverse samples, including clinical populations with more severe anhedonic symptoms, will therefore be needed to test whether the focal anhedonia-related effects observed here are reliable and generalisable. Repeated-measures or longitudinal designs could further clarify the stability of these effects over time, although the primary question addressed in the present study did not require longitudinal data.

The present analyses also collapsed across happy, sad, and neutral excerpts, which may have obscured valence-specific patterns of neural synchrony. This is relevant because recent work suggests that affective valence may shape inter-individual neural asymmetrically, with negative affect associated with more similar default-network connectivity profiles and positive affect associated with more idiosyncratic profiles during social memory consolidation (Iyer et al., 2024). Future analyses separating musical excerpts by affective valence may therefore better test whether anhedonia is associated with altered synchrony for positive versus negative music.

Second, although ROI analyses were corrected within masks, correction was not applied across the full set of masks and models. The right amygdala and left insula findings should therefore be treated as exploratory until replicated in an independent sample. In addition, ROI analyses were implemented by extracting pairwise ISC values from existing whole-brain ISC outputs within each mask, rather than rerunning ISC constrained to each ROI. The resulting maps should therefore be interpreted as voxel-wise summaries within predefined ROIs rather than independent ROI-specific ISC analyses.

More broadly, ISC analyses assume consistent voxel-wise anatomical and functional alignment across participants (Nastase et al., 2019), an assumption that may be violated in some higher-order and affective regions where inter-individual variability is especially pronounced (Mueller et al., 2013). Conventional group-level ISC is also relatively insensitive to idiosyncratic but meaningful neural responses that are not temporally synchronised across participants. Pairwise ISC and IS-RSA partly mitigate this limitation by preserving variability in response similarity across participant pairs, rather than collapsing directly to a single group-level synchrony estimate. However, these analyses still depend on sufficient temporal and spatial correspondence between participants; meaningful affective responses that occur at slightly different times or locations across individuals may still produce low ISC. These considerations warrant caution when interpreting weak or null ISC effects in affective regions.

### Conclusion

This study demonstrated that naturalistic music listening elicited robust inter-subject neural synchrony, particularly within auditory and sensorimotor regions. However, shared neural activity showed only limited correspondence with similarity in subjective emotional ratings, suggesting that common perceptual responses do not necessarily translate into shared affective experience. Continuous anhedonia IS-RSA provided limited evidence for systematic associations between MASQ-AD scores and neural synchrony, with most targeted ROI masks and the exploratory whole-brain analysis showing no FDR-corrected effects. A focal negative association in the right amygdala survived within-mask FDR correction, suggesting that anhedonia-related variation in neural synchrony may be region-specific.

Together, these findings support the view that music listening involves both shared perceptual-sensorimotor processing and more individualised emotional appraisal. Future studies employing advanced analytical approaches and larger, more diverse samples will be needed to clarify how shared and individualised neural responses during naturalistic emotional experiences relate to affective traits such as anhedonia.

## Acknowledgements

The authors acknowledge support from the National Institute for Health and Care Research (NIHR) Maudsley Biomedical Research Centre at South London and Maudsley NHS Foundation Trust and King’s College London. The views expressed are those of the authors and not necessarily those of the NIHR, NHS, or the Department of Health and Social Care.

## Supplementary Materials

